# Bayesian inference of agent-based models: a tool for studying kidney branching morphogenesis

**DOI:** 10.1101/096032

**Authors:** Ben Lambert, Adam L. MacLean, Alexander G. Fletcher, Alexander N. Combes, Melissa H. Little, Helen M. Byrne

## Abstract

The adult mammalian kidney has a complex, highly-branched collecting duct epithelium that arises as a ureteric bud sidebranch from an epithelial tube known as the nephric duct. Subsequent branching of the ureteric bud to form the collecting duct tree is regulated by subcellular interactions between the epithelium and a population of mesenchymal cells that surround the tips of outgrowing branches. The mesenchymal cells produce glial cell-line derived neurotrophic factor (GDNF), that binds with RET receptors on the surface of the epithelial cells to stimulate several subcellular pathways in the epithelium. Such interactions are known to be a prerequisite for normal branching development, although competing theories exist for their role in morphogenesis. Here we introduce the first agent-based model of *ex vivo* kidney uretic branching. Through comparison with experimental data, we show that growth factor-regulated growth mechanisms can explain early epithelial cell branching, but only if epithelial cell division depends in a switch-like way on the local growth factor concentration; cell division occurring only if the driving growth factor level exceeds a threshold. We also show how a recently-developed method, “Approximate Approximate Bayesian Computation”, can be used to infer key model parameters, and reveal the dependency between the parameters controlling a growth factor-dependent growth switch. These results are consistent with a requirement for signals controlling proliferation and chemotaxis, both of which are previously identified roles for GDNF.

**Author Summary:** A number of important congenital disorders arise due to incomplete development of the mammalian kidney. Elucidating the cause of these conditions requires an understanding of the mechanisms that contribute to kidney morphogenesis. Whilst experimental work has suggested several candidate mechanisms, their importance is still not well understood. Here we develop a computational model of kidney morphogenesis at the individual cell level to compare these different hypotheses. Guided by existing experimental evidence we propose that a generic growth factor, that we term “GDNF”, produced from the mesenchyme surrounding the epithelium, can drive a number of cellular responses. Simulations of our agent-based model reveal that diffusion of GDNF, coupled with GDNF-stimulated epithelial cell division, can generate the branching patterns seen in *ex vivo* kidney explant experiments. We also find that branching depends on the sensitivity of cell proliferation to changes in GDNF levels. In particular our model only generates realistic branching when there is significant variation in GDNF levels along the boundary of the epithelium, and most cells divide only if the local concentration of GDNF exceeds a threshold value. We conclude that feedback between mesenchymal cells that produce GDNF, and epithelial cells that consume it, is vital for normal kidney organogenesis.

## 1 Introduction

The kidney is a complex organ with a highly branched structure. Its primary function is to filter urea and other waste products from the blood and metabolic system. Kidney and urinary tract congenital disorders are amongst the most common birth defects, with many of these conditions being caused by incomplete branching during development (Airik & Kispert, 2007). To understand how such disorders arise, we must first address fundamental questions about the developing kidney, including how branching is initiated, how it is regulated, and how cessation of branching is controlled. These processes are complex and, as such, biological experiments that aim to answer these questions are technically difficult to conduct. Furthermore, each experiment typically investigates only a single facet of kidney morphogenesis. Mathematical and computational models can assist in this endeavour by integrating multiple biological hypotheses via a well-defined set of assumptions. Simulations of the resulting models can be performed to test alternative hypotheses about the mechanisms that regulate kidney morphogenesis, and generate new predictions that can be tested experimentally (Cebrian *et al.*, 2014; Combes, 2015; Murray *et al.*, 1983; Packard *et al.*, 2013; Short *et al.*, 2014; Zubkov *et al.*, 2015).

Nephrons are the primary functional units of a kidney. A fully developed human kidney has between 0.2 and 1.8 million nephrons (Hughson *et al.*, 2003), that are connected by a system of collecting ducts to the ureter and bladder (Costantini, 2006). Most of the organ is formed during embryonic development, beginning five weeks post-gestation in humans and embryonic day 10.5 (E10.5) in mice (Carlson, 2013; Cebrian *et al.*, 2004). An outgrowth of epithelial cells from the nephric duct leads to the formation of the ureteric bud epithelium (Little, 2015). As the ureteric bud grows and branches, a cloud of mesenchymal progenitor cells form caps at the tips, and eventually differentiate into nephrons.

In addition to their role as nephron precursors, mesenchymal cells release cytokines that influence the growth and branching of epithelial cells as they form the collecting tree structure. Glial cell line-derived neurotrophic factor (GDNF) is one such growth factor. GDNF is expressed in the metanephric mesenchyme by E11.5, and is thought to diffuse across to the epithelial cell layer, where its binding to cell surface RET receptors transduces signals that are essential for morphogenesis (Durbec *et al.*, 1996). GDNF is known to be a chemoattractant, and also to stimulate the outgrowth of epithelial cells (Little & McMahon, 2012). However, there is no consensus about whether these mechanisms are sufficient to generate normal branching, or whether additional chemical or mechanical mechanisms are needed. For example, GDNF-independent signals involving members of the fibroblast growth factor family and Activin are known to affect branching (Maeshima *et al.*, 2007; Michos *et al.*, 2010; Miyazaki *et al.*, 2000; Qiao *et al.*, 2001; Tee *et al.*, 2010, 2013). As the roles of GDNF and other growth factors remain to be fully elucidated, in this paper we focus on a single generic growth factor which we term “GDNF” and attribute to it actions and effects that we acknowledge are likely caused by a combination of different factors. Although the mechanical forces between epithelium and mesenchyme may influence branching patterns, they are not a prerequisite for branching. Indeed experiments performed by Qiao *et al.* (1999) have revealed that cell–cell contact between mesenchyme and epithelium is not required for branching; exposure of epithelial cells to soluble factors derived from metanephric mesenchyme is sufficient. While it is possible that mechanical forces between the epithelial cells may be an important mechanism for branching, to our knowledge no experiments have yet directly investigated this hypothesis.

Branching morphogenesis is a characteristic feature of many mammalian organs including the kidney, lung, vasculature, the saliva and mammary glands, and in limb development (Affolter *et al.*, 2009). In each of these tissues, branched structures arise due to repetition of three cellular motifs:*bud formation* followed by *bud extension*, and *bud splitting*, with these processes being facilitated by regulation at the tips and stalks of individual buds. Similar pathways and network topologies perform this regulation in vastly different systems; for example, fibroblast growth factor signalling in combination with Delta/Notch signals regulate branching of both trachea and vertebrate vasculature (Affolter *et al.*, 2009; Ochoa-Espinosa & Affolter, 2012). Several alternative theories have been proposed to explain how such signalling generates branched structures. For example, Turing’s reaction-diffusion mechanism can give rise to branched structures (Menshykau & Iber, 2013). Mechanical theories have also been proposed (Varner & Nelson, 2014). To distinguish between competing theories, simple systems are needed, whose variability can be controlled. Explant models of branching have a valuable role to play here as they can recapitulate branching morphogenesis, and are amenable to image analyses that are difficult or impossible to perform *in vivo* (Basson *et al.*, 2006; Serls *et al.*, 2005; Watanabe & Costantini, 2004). Additionally, it is possible not only to monitor the structure of an explant as it evolves over time but also to determine for example how these dynamics change when the composition of the culture medium that bathes the explant is altered.

A variety of mathematical models have been developed to study different aspects of branching morphogenesis, including physical cellular processes, such as proliferation and migration (Hirashima *et al.*, 2009), the underlying molecular processes (Zubkov *et al.*, 2015), or a combination of the two (Adivarahan *et al.*, 2013; Clément & Mauroy, 2014; Menshykau & Iber, 2013). A recent study used ordinary differential equations to describe how the dynamics of the growing epithelial cell populations at the tips of branches are regulated by mesenchymal cells (Zubkov *et al.*, 2015). Comparison of simulation results and experimental data revealed that the mathemematical model could recapitulate the observed dynamics for the ratio of epithelial to mesenchymal cells at the branch tips. In other studies a Turing system (Turing, 1952) involving the diffusion of the secreted ligands GDNF (produced by mesenchyme) and Wnt11 (produced by the epithelium in response to the presence of GDNF), as well as RET receptors, is proposed to be the primary driver of branching (Clément & Mauroy, 2014; Menshykau & Iber, 2013). Predictions generated by these models include the need for cooperative receptor-ligand binding (Menshykau & Iber, 2013), and a mechanism for self avoidance (so that growing tips do not come into contact) (Clément & Mauroy, 2014), which recent experiments suggest may be orchestrated by signalling via BMP7 that is produced by neighbouring branches (Davies *et al.*, 2014).

Whilst a few models of developmental processes consider individual cell behaviour (Fletcher *et al.*, 2014; McLennan *et al.*, 2012, 2015), the majority treats cell populations as a continuum (Clément & Mauroy, 2014; Menshykau & Iber, 2013; Scialdone *et al.*, 2013; Zubkov *et al.*, 2015). While these models have generated valuable insight into different aspects of organ development, they are limited in their ability to investigate the influence of cellular and subcellular mechanisms. Agent-based frameworks that model individual cell behaviour can offer significant advantages here.

In a biological context, agent-based models treat cells as individual agents, whose behaviour is specified by a predefined set of rules that can be deterministic and/or stochastic. First developed to study the dynamics of replication (von Neumann, 1966), simple agent-based models such as cellular automata (CA), and generalisations of these such as cellular Potts and hybrid models are now used widely to study a range of biological systems that include biochemical reaction networks, stem cell proliferation and differentiation, tumour angiogenesis and metastasis (Alarcón *et al.*, 2003; Gerlee & Anderson, 2015; Macklin *et al.*, 2012; Roeder *et al.*, 2006; Scott *et al.*, 2014). Such “lattice-based” models restrict agents to sites on a fixed lattice. As such they are typically simpler in design and faster to simulate than alternative off-lattice models, which place fewer restrictions on the movement of cells (or agents) (Bentley *et al.*, 2009; Pathmanathan *et al.*, 2009; Perfahl *et al.*, 2016).

In this paper we investigate a series of experiments performed by Watanabe & Costantini (2004) in which kidneys from mouse embryos were grown in culture. We develop an agent-based framework to model the growth of these explants, and focus on interactions between epithelial cells and growth factors (referred to as “GDNF”) present in the culture medium. Since the experimental data we have available is limited in detail we formulate an idealised model that contains the minimal number of assumptions necessary to recapitulate key branching features. Whilst the model does not explicitly account for mesenchyme cells, local levels of the generic growth factor, which we term GDNF, serve as a proxy for their influence. We use a CA approach which does not require specific assumptions about the nature of cell-cell forces that cannot (as yet) be experimentally verified. In more detail, our agent-based epithelial cells reside on a regular, two-dimensional grid, their rates of migration and proliferation being regulated by (and, in turn, regulating) the local distribution of GDNF.

Typically the parameters of computational models cannot be directly measured; they must be inferred from experimental data. Inference can be comparatively expensive in agent-based models, with the stochastic nature of the models requiring multiple simulations for each choice of parameter values. This cost can preclude parameter estimation even with state-of-the-art techniques, such as approximate Bayesian computation (ABC) (Johnson *et al.*, 2014; Liepe *et al.*, 2014; Toni *et al.*, 2009). ABC compares simulations from a model with experimental data, and – using statistics that summarise the system behaviour – accepts simulations (and the parameters that generated them), if the statistics for model and data lie within an acceptable distance threshold (see for example, Johnston *et al.* (2014); Vo *et al.* (2015)). ABC relies on the ability to simulate a model relatively quickly (typically faster than one second per one stochastic realisation of the model). In situations where this condition is not met, new methods are required for parameter inference. One such method is “Approximate Approximate Bayesian Computation” (AABC) (Buzbas & Rosenberg, 2015). With AABC actual model simulations are used to generate pseudo-replicates of the model that can be compared with the data. Since these pseudo-data are generated in a fraction of the time needed to perform real simulations, this method can exhibit significant speed up compared to ABC. Our agent-based CA model takes approximately a minute to generate a single simulation on a desktop computer, and hence we chose to use AABC for parameter inference.

We analyse the model first by direct simulation across a range of parameter regimes, in each case comparing model outputs and experimental data. We next use AABC to quantitatively fit the model to data, and reveal its dependence on key parameters. In doing so, we show how AABC can be used to integrate the agent-based model (ABM) and the experimental data to answer the following questions:

1. Does the mechanism of GDNF-dependent cellular growth (that is, where cell proliferation rates increase with increases in the local concentration of GDNF) lead kidney explants to develop branches as seen *ex vivo*?
2. Which characteristics of GDNF-mediated mechanisms are necessary to generate these branches?
3. How sensitive is the model to changes in those parameters that influence branching?

We now briefly outline the structure of the paper. In Section 2.1 we introduce the experimental data that we use in this study, and explain the details of how they are processed to yield summary statistics that represent key features of explant branching. In Section 2.2 we describe our CA model and, in doing so, explain how individual cell behaviour depends on a field of GDNF that diffuses through the simulation domain. In Section 2.3 we explain how we implemented the AABC method, and then use it to investigate the factors that affect explant branching. In Section 3.1 we demonstrate how our model is able to recapitulate the branching of kidney explants, and in Section 3.2 we use direct simulation to determine how branching of our model explants depends on specific biological processes. Finally in Section 3.3 we use AABC to quantitatively investigate how explant branching depends on three model parameters that quantify the rate of cell motility and cell division.

## 2 Materials and Methods

### 2.1 Kidney explant data experimental data and image processing

Watanabe & Costantini (2004) have developed an *ex vivo* assay to study epithelial branching in murine embryonic development. Kidneys were dissected from E11.5 embryos from a Hoxb7/EGFP transgenic line expressing green fluorescent protein (GFP) throughout the nephric duct and the ureteric bud (Srinivas *et al.*, 1999). Kidneys were cultured in fetal bovine serum and imaged every 30 minutes over a period of 96 hours, generating movies of the developing explants.

We processed videos of the experimental data (three explants in total) by extracting the area of the epithelial mass and the number of branch points in each frame. All image processing and analyses were performed using Matlab (MathWorks, Natick, MA). We started by extracting the bulk epithelial mass and then used this to calculate the medial axis skeleton (see Figs. 1, 3): this can be derived as the loci of centres of bi-tangent circles that fit entirely within the epithelial region (Lee, 1982). We then counted the number of branch points on each skeleton and used this as a measure of the amount of branching that had occurred. Calculation of the medial axis skeleton and counting of branching points were carried out using Matlab’s “bwmorph” function, together with a third-party Matlab package (available from “https://uk.mathworks.com/matlabcentral/fileexchange/11123-better-skeletonization”). While this summary statistic captures the aggregate number of branches, it does not explain other features such as whether the branches are primary or secondary, nor does it provide information about the tip-to-stalk ratio (Bush *et al.*, 2014; Little, 2015). However we consider the collection and analysis of such detailed summary statistics to be overly sophisticated for comparison with our CA model and, hence, we postpone this for future work.

**Figure 1.**
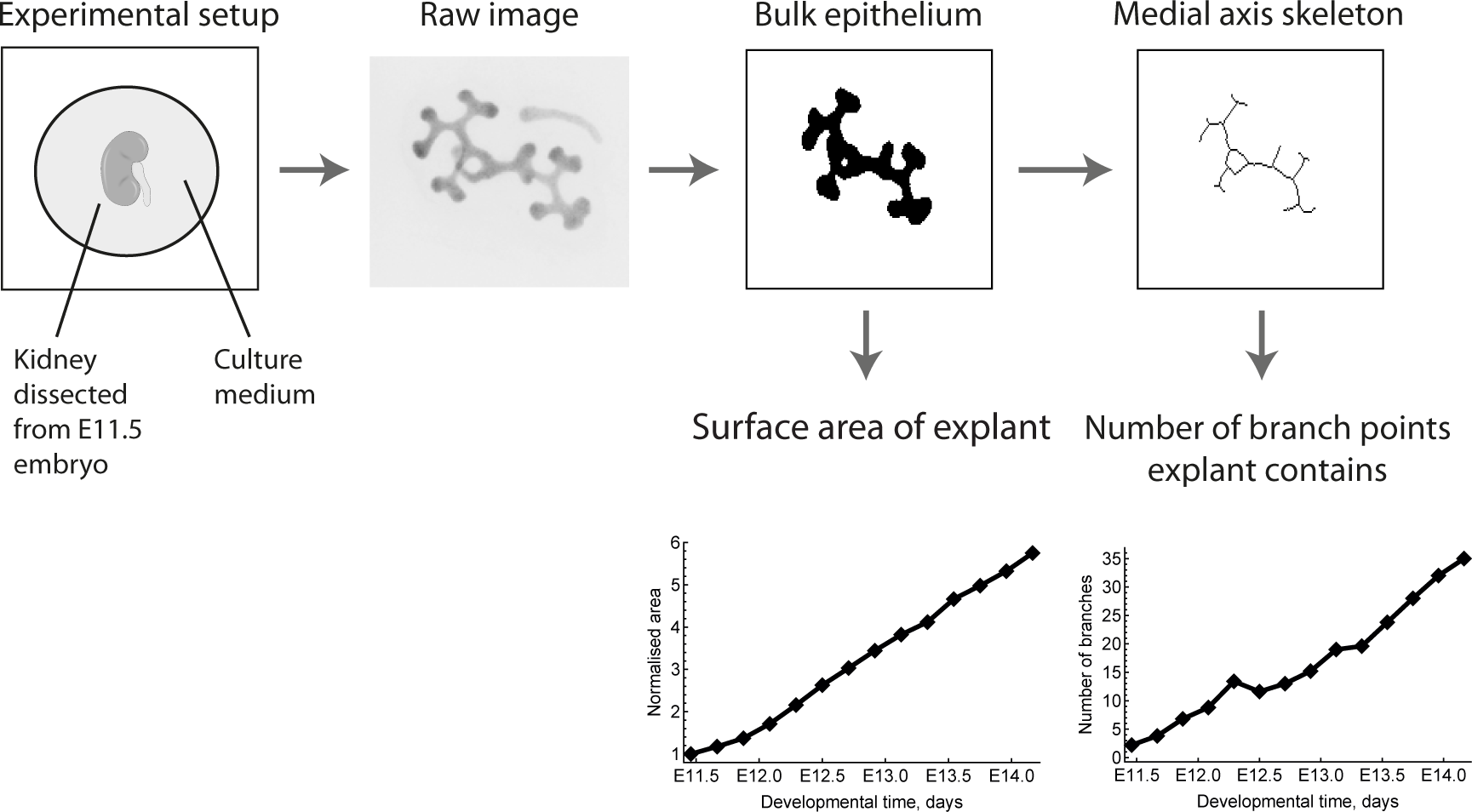
Workflow used to extract data from the explant images. The same process is applied to simulation results generated from the CA model. The raw image is reproduced with permission from Watanabe & Costantini (2004).

For comparison with our model simulations we also monitored the area of the bulk epithelium as a measure of the change in the number of cells over time. To facilitate comparison of the data from each experiment we normalised the area of each epithelial mass at time *t* by its initial mass (at *t* = 0).

For each CA simulation (described below) we recorded the positions of all epithelial cells over time. We then applied the same image processing methods to the CA data as were applied to the experimental data, in order to generate comparable summary statistics for branching and cell numbers.

### 2.2 Cellular automaton model development

Our model comprises a two-dimensional cellular automaton (CA) model in which the behaviour of individual epithelial cells (i.e. their speed and direction, their rate of division and consumption of GDNF) depends on the locations of nearby cells and the local concentration of GDNF. Cell-cell interactions are governed by rules chosen to replicate observed behaviour. Our model is admittedly an idealisation of the biological processes that underpin kidney morphogenesis. In the absence of suitable experimental data, our model does not explicitly include mesenchyme cells even though they likely surround the explanted epithelium (they are invisible in the experimental images). Similarly our model does not account for a GDNF-Wnt11 positive feedback loop: GDNF secreted by mesenchymal cells binds to RET receptors on epithelial cells and stimulates them to produce Wnt11 which binds to mesenchymal cells, stimulating further uptake of GDNF (Majumdar *et al.*, 2003). However since the available experimental data provides limited information we chose to formulate a simple model that could reproduce key features of kidney morphogenesis.

The epithelial cells occupy a square domain which is discretised into *N × N* equally-spaced grid points. Each lattice site is occupied by either an epithelial cell or extracellular matrix (ECM). At *t* = 0, a mass of epithelial cells is introduced towards the centre of the domain. Its shape and size (i.e. the number of cells) are chosen to resemble those from the initial images of kidney explants from Watanabe & Costantini (2004). All other sites are occupied by ECM at *t* = 0.

#### GDNF field

We assume that GDNF binds to receptors on the outer membrane of epithelial cells at rate *K_G_*, and diffuses from the boundaries of the grid, where it is maintained at a constant GDNF concentration, *G_∞_*. This implicitly represents a far-field approximation of the experimental conditions, where the ECM area is large, and the local concentration of GDNF is continuously replenished. These conditions are chosen to mimic the effects of GDNF produced by mesenchymal cells (that are likely present) as well as other growth factors that are present in the culture medium. Since the time scale for the diffusion of GDNF across the length of a cell diameter (seconds) is much shorter than the time scale for cell division (hours), we use a quasi-steady-state 2D reaction-diffusion-equation (with Dirichlet boundary conditions) to model the distribution of GDNF, *G* (*x, y, t*),

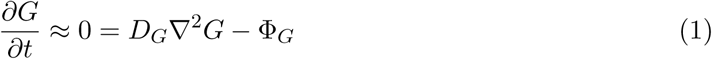

where *D_G_* denotes the assumed-constant diffusion coefficient for GDNF, and Φ_*G*_ is the local rate of GDNF consumption,

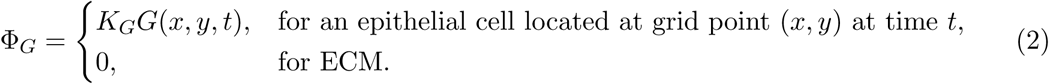

In eqn. (2) we assume, for simplicity, that the rate of GDNF consumption depends linearly on the concentration of substrate (with rate parameter *K_G_*). In what follows, it is convenient to recast eqn. (1) in terms of a dimensionless GDNF concentration 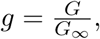 and non-dimensional spatial parameters, 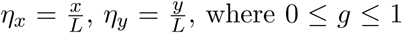 is the GDNF concentration as a ratio of that at the boundaries, and 0 ≤ *η_x,y_* ≤ 1 are spatial coordinates as a fraction of the simulation domain size. Eqn. (1) can then be written as,

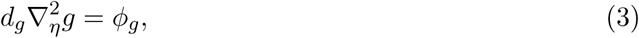

where 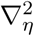 is the Laplacian with respect to the non-dimensional spatial coordinates, 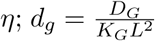 is the non-dimensional diffusion coefficient; and 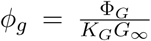 is the non-dimensional GDNF uptake term. In these coordinates the boundary conditions are specified as,

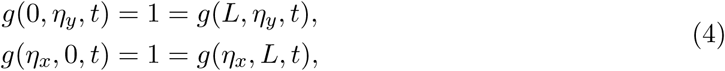

and the local rate of GDNF consumption is given by,

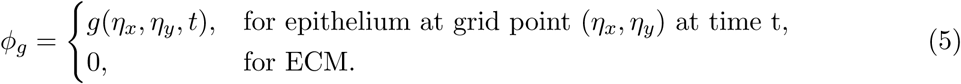

In the absence of experimental estimates for the parameters *D_G_* and *K_G_*, preliminary numerical experiments revealed that fixing *d_g_* = 0.006 lead to sufficient variation in GDNF across the spatial domain for branching to occur. (See Table S1 for a summary of the parameter values used in each simulation.)

At the end of each time step (see “Cell based rules” section) we solved eqns. (3), (4) and (5) using the method of explicit finite differences (implemented in Matlab). Both the CA model and the finite difference scheme are implemented on the same discrete grid, so the local GDNF level used in the update rules is its value at the grid point where the cell is located. The GDNF field affects the movement and behaviour of epithelial cells, while their location and rates of GDNF uptake affect the evolution of the GDNF field (Fig. 2).

**Figure 2.**
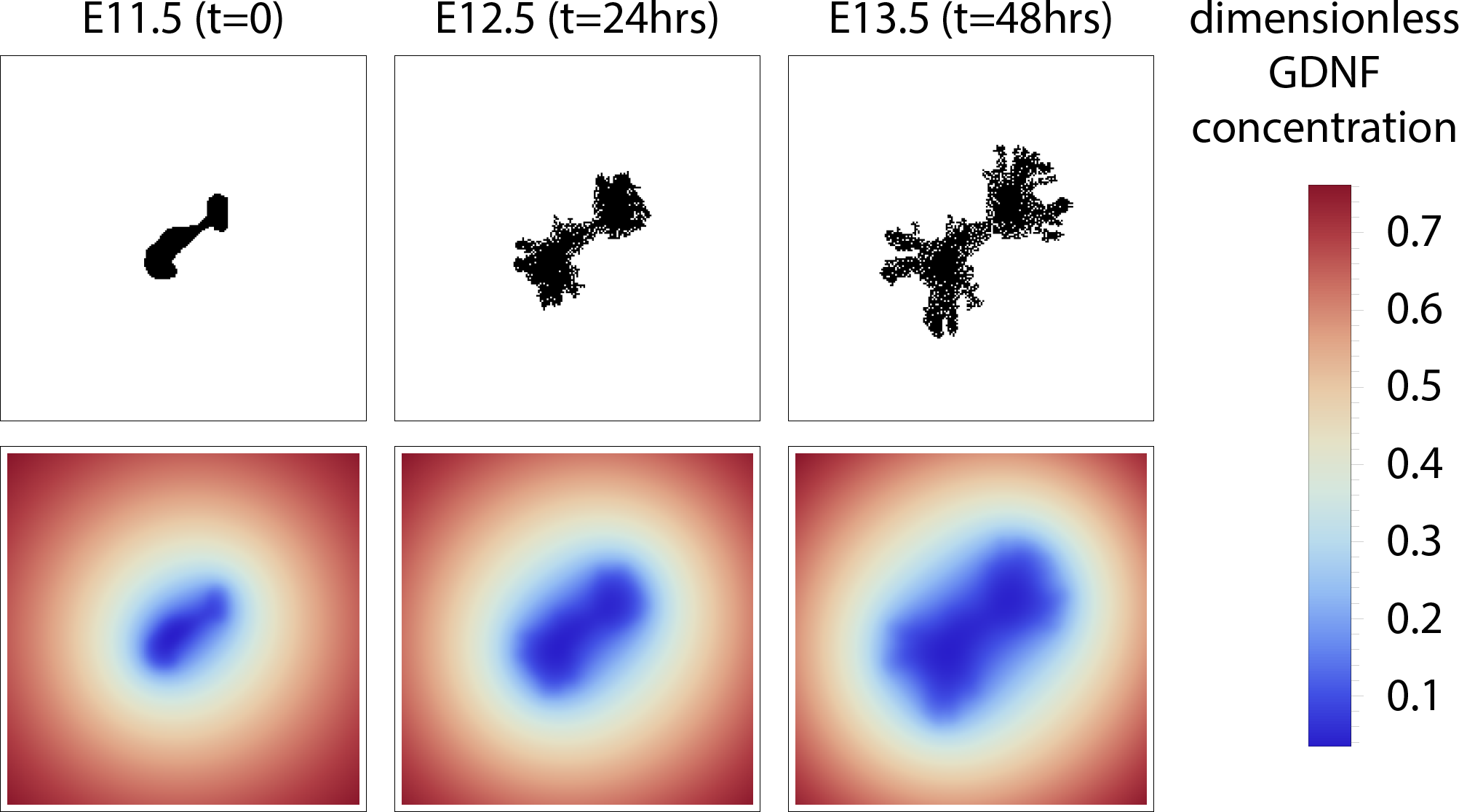
A series of plots from a typical CA simulation illustrating the interaction between epithelial cell positions (top) and GDNF level (bottom). Note: the GDNF level at the edges is not *g_1_* = 1 because the panels only show the central 140*×*140 grid points of the domain (itself of size 400*×*400). The parameter values used to generate these plots are shown in Table S1.

#### Cell-based rules

Simple rules are used to determine whether individual cells move or divide (see Algorithm 1, and Table S1 for the parameter values used); cell death is assumed to be negligible in line with previous experimental results (Hartman *et al.*, 2007). The algorithm begins by solving for the steady state solution of the reaction-diffusion equation (as described above) for the current epithelial cell locations. Using a randomly-permuted list of cell indices, each cell is then visited in turn, and updated. Updates proceed as follows. If a cell has empty neighbouring sites then we determine whether to propose a movement or division event into one of the empty locations; the local GDNF concentration determines whether the proposed action occurs and, if so, the empty site at which the action is carried out.

During this process we establish whether any of the 4 adjacent sites in the von Neumann neighbourhood (up, down, left, right) are empty (no epithelium present). If this is the case then we select a move with probability *p_move_*, or a cell division with probability (1 − *p_move_*). The probability *p_move_* is independent of GDNF, and is used simply to propose an action (that may not be undertaken). If an action is selected and feasible, our cell-based rules dictate whether it is executed. Moves are always carried out; whether cell division occurs depends on local levels of GDNF. We refer to this process as “GDNF-stimulated cell division”. In particular, the probability *p_cd_* that a cell at location (*η_x_*, *η_y_*) at time *t* divides is given by,

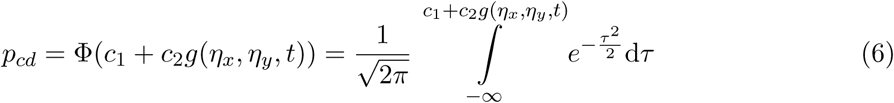

where Φ(.) is the standard normal cumulative distribution function. This functional form allows a switch-like behaviour for cell division. In eqn. (6) the parameter *c*_1_ controls the location of the switch, while the parameter *c*_2_ determines its sensitivity to the local GDNF level, *g*(*x, y, t*). In most simulations we fix *c*_1_ *<* 0 and *c*_2_ *>* 0, so that in the absence of GDNF (*g* = 0) by default epithelial cells do not divide.

If a move or cell division event is to occur, we must decide which empty neighbouring grid point will be occupied by the result of the action. This choice (if there is more than one empty grid point) is biased by GDNF levels at the neighbouring sites. For moves this process represents “chemotaxis” and for cell divisions it represents “anisotropic cell division”. In either case the probability *p_i_* of selecting an empty neighbouring grid point 1 ≤ *i* ≤ *K* is given by,

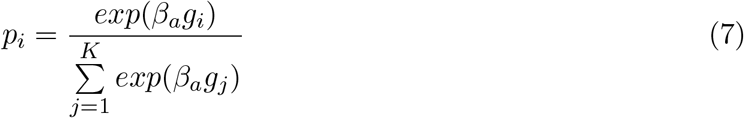

where *a* ∊ {move, cell-division} specifies the action type, 1 ≤ *K* ≤ 4 is the number of empty neighbours, and *g_i_* is the GDNF concentration at site *i*. In eqn. (7) the non-negative parameter *β_a_* controls the sensitivity of the selection to GDNF concentration, which can differ for chemotaxis and anisotropic cell division.

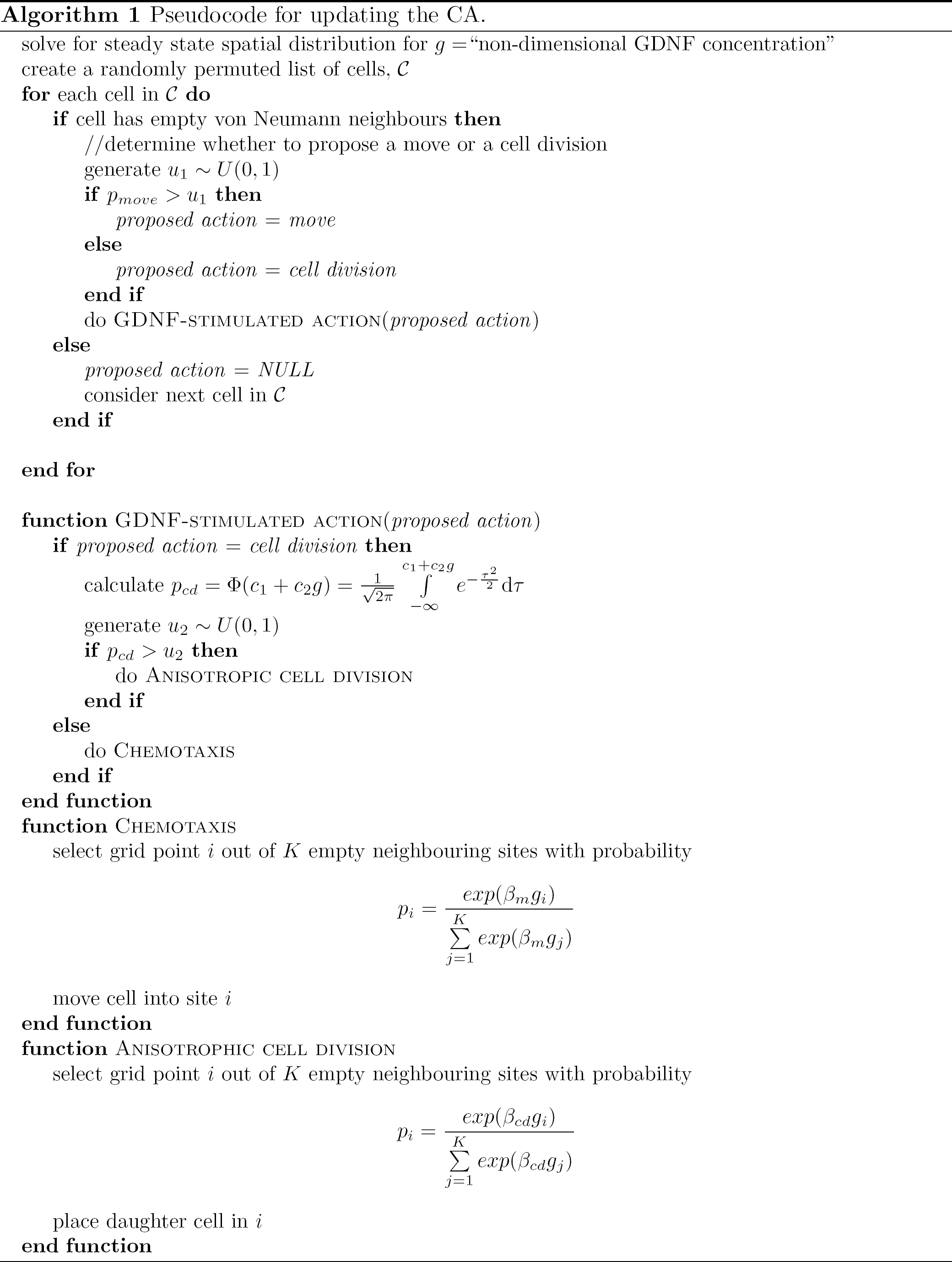

### 2.3 Parameter inference for agent-based models

Approximate Bayesian computation (ABC) allows parameter inference and model selection using distance-based criteria to compare simulations and data (Sunnåker *et al.*, 2013). A distance metric is introduced (e.g. Euclidean or Manhattan) and simulations are rejected if they yield values for a (set of) chosen statistic(s) that exceed a threshold value. ABC is appropriate when a likelihood function is difficult or intractable to calculate. It has been applied to a wide range of problems in systems biology (Beaumont *et al.*, 2002, 2010; Csilléry *et al.*, 2010a). A prerequisite of ABC is that simulating a dataset from the model must be relatively inexpensive so that it is possible to adequately sample from the posterior parameter space. This is the case whether performing simple ABC rejection or more sophisticated procedures, such as sequential Monte Carlo ABC (Toni *et al.*, 2009). A simulation time for a single dataset of *O* (*seconds*) or greater is thus typically prohibitive for performing ABC.

A new algorithm named “approximate approximate Bayesian computation” (AABC) has been designed to resolve this problem (Buzbas & Rosenberg, 2015). By replacing true model simulations with realisations of a suitably-tuned statistical model, AABC enables parameter inference for a class of models that previously presented large (or insurmountable) computational challenges. Agent-based models, such as our CA model, represent such a class, that have with few notable exceptions (Johnston *et al.*, 2014; Jones *et al.*, 2015; Sottoriva & Tavare, 2010), evaded approximate Bayesian inference.

Here we use AABC to infer parameters of the CA model that yield behaviour consistent with the average summary statistics extracted from dynamic experimental data for kidney explants generated by Watanabe & Costantini (2004). These summary statistics (see section 2.1 above) measure two facets of the evolving explants: the number of branches, and the rate of cell proliferation. By comparing summary statistics from the experimental data with equivalent statistics from our simulations (as detailed below) we investigate the sensitivity of explant growth patterns to variation in three model parameters *c*_1_, *c*_2_, and *p_move_*. We specify uniform priors for each of these parameters with the following bounds:*c*_1_ ∊ [*−*20*, −*40], *c*_2_ ∊ [40, 280] and *p_move_* ∊ [0, 1]. All other parameters are held fixed (as per Table S1).

Specifically the algorithm implements the following steps:

1. Simulate the CA model, and accept a subset of particles with parameter sets *θ_i_* = (*c*_1*i*_, *c*_2*i*_, *p_movei_*) and corresponding datasets *x_i_* = (*x*_1*i*_, *x*_2*i*_), *i* ∊ (1, 2*, …, m*), where *m* is the number of simulations (each with a unique set of parameters sampled from the priors) and *N* = 2 is the number of replicates per parameter set. For our application, each dataset *x* consists of data for the normalised area and the number of branch points at *t* = 10, 20, 30 hours after E11.5.
2. Sample a new set of parameter values, *θ\**, from the prior.
3. Calculate the weights, *ω_i_*, using an Epanechnikov kernel (Buzbas & Rosenberg, 2015):

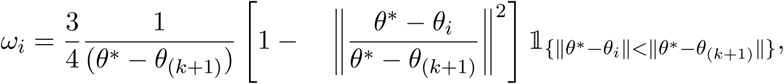

where 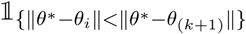 is an indicator function with value 1 for the *k* parameter values *θ* = *θ*_1_*, θ*_2_, …, *θ_k_* with the shortest Euclidean distance from *θ\**, and 0 otherwise (from *θ_k+1_* onwards).
4. Select (*θ_i_*, *x_i_*), *i* ∊ (1, 2*, …, k*), for which *ω_i_* > 0.
5. Draw a sample *ϕ* – used to specify data resampling probabilities – from a Dirichlet distribution parameterised by *ω_i_*, *i* ∊ (1, 2*, …, k*).
6. Simulate a new dataset x* of area and branching time point data (of size *N* = 2 replicates) by: (*i*) resampling datapoints from *x_i_* with probabilities set by *ϕ*, and (*ii*) assuming that each replicate is equally probable.
7. Calculate the Euclidean distance between the real and simulated datasets, and add *θ\** to the posterior, iff ||x_*i*_ − x^*^|| < ∊. We choose *∊* such that 5% of simulations are accepted as posterior samples.
8. Repeat steps (2)-(7) until convergence in the approximate posterior distribution is reached.

We perform Step (1) of the algorithm in Matlab (MathWorks, Natick, MA), and steps (2)-(7) in Julia (v0.3.5, julialang.org).

## 3 Results

### 3.1 GDNF-directed cellular proliferation can explain the branching patterns observed when kidney explants are cultured *ex vivo*

Simulations of the CA model reveal that it can generate branching patterns similar to those identified from *ex vivo* kidney explant data of Watanabe & Costantini (2004). In Fig. 3 we compare results from a typical simulation with experimental data collected at four time points. The CA model recapitulates notable features of branching: branching at both ends of the buds, followed by secondary branching at the branch tips. Our simulated explants also often produce branching events in which three or more branches emerge from a single tip, events which are observed in the developing kidney (Menshykau & Iber, 2013).

**Figure 3.**
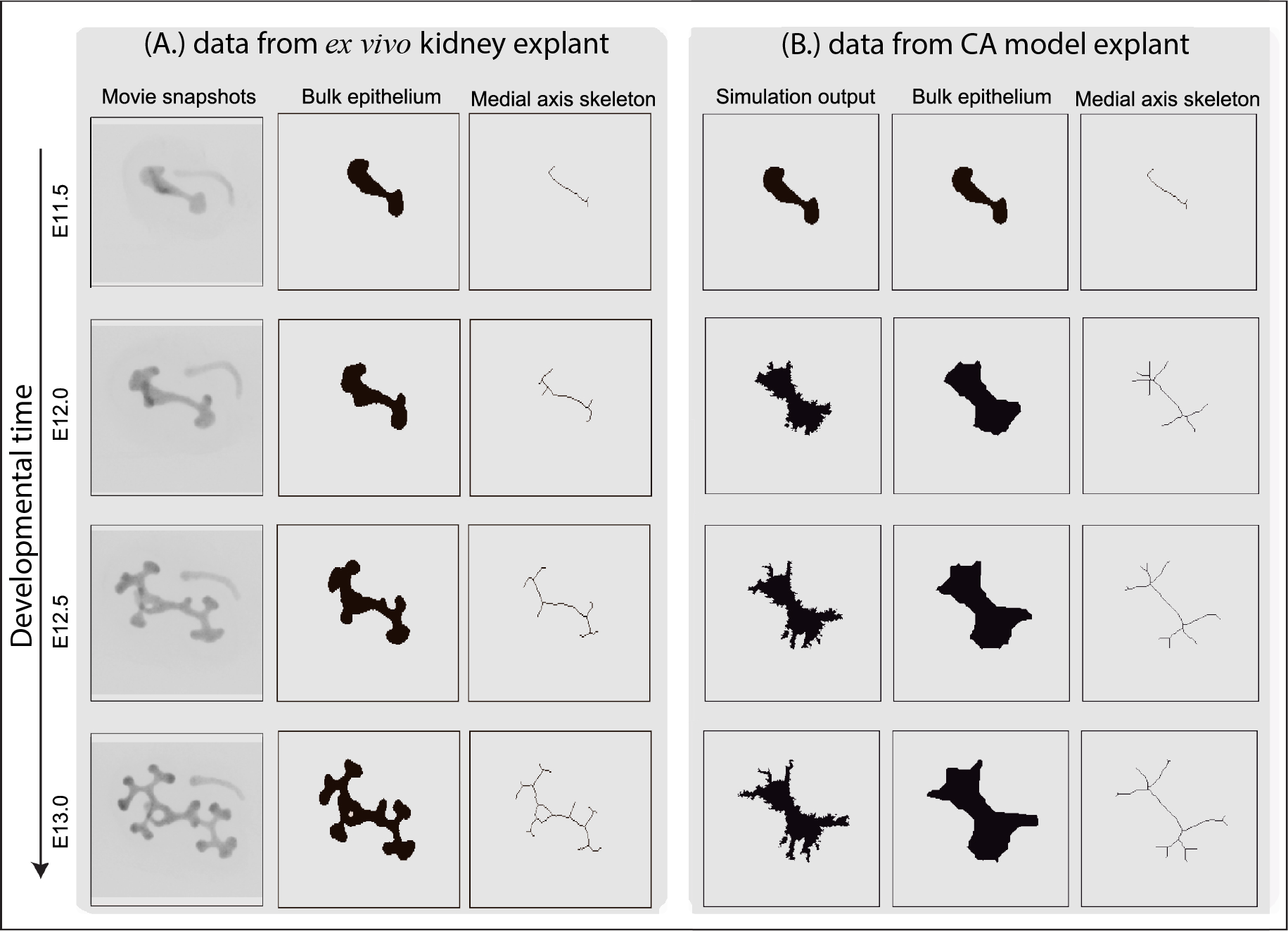
Comparison of branching patterns associated with (A.) *ex vivo* data and (B.) typical simulations of the CA model. Image processing techniques (see section 2.1) are used first to extract the bulk shape of the epithelial cells (middle panel) and then to generate the medial axis skeleton (right panel), which characterises the shape of the growing explant. The parameter values used to generate these simulation results are shown in Table S1.

### 3.2 Finding a minimal set of sub-cellular mechanisms necessary to generate branching

We also used our model to investigate the contribution of cell proliferation, anisotropic cell division and chemotaxis to branching (Table 1 and Figs. 4 and 5). Each mechanism is regulated by local levels of GDNF: the rate of cell division increases with local levels of GDNF in a sigmoidal manner (see Fig. 6); regarding chemotaxis, cells are more likely to migrate up local GDNF gradients; for anisotropic cell division (ACD), daughter cells are located preferentially in sites with higher levels of GDNF. For both chemotaxis and ACD we acknowledge that our model is an idealisation of the biological processes involved. For example RET-dependent movement (Riccio *et al.*, 2016) and luminal mitosis (Packard *et al.*, 2013) are not considered explicitly. Even so, our model captures some of the features of these processes and, hence, can be used to determine their relative contributions for branching.

**Figure 4.**
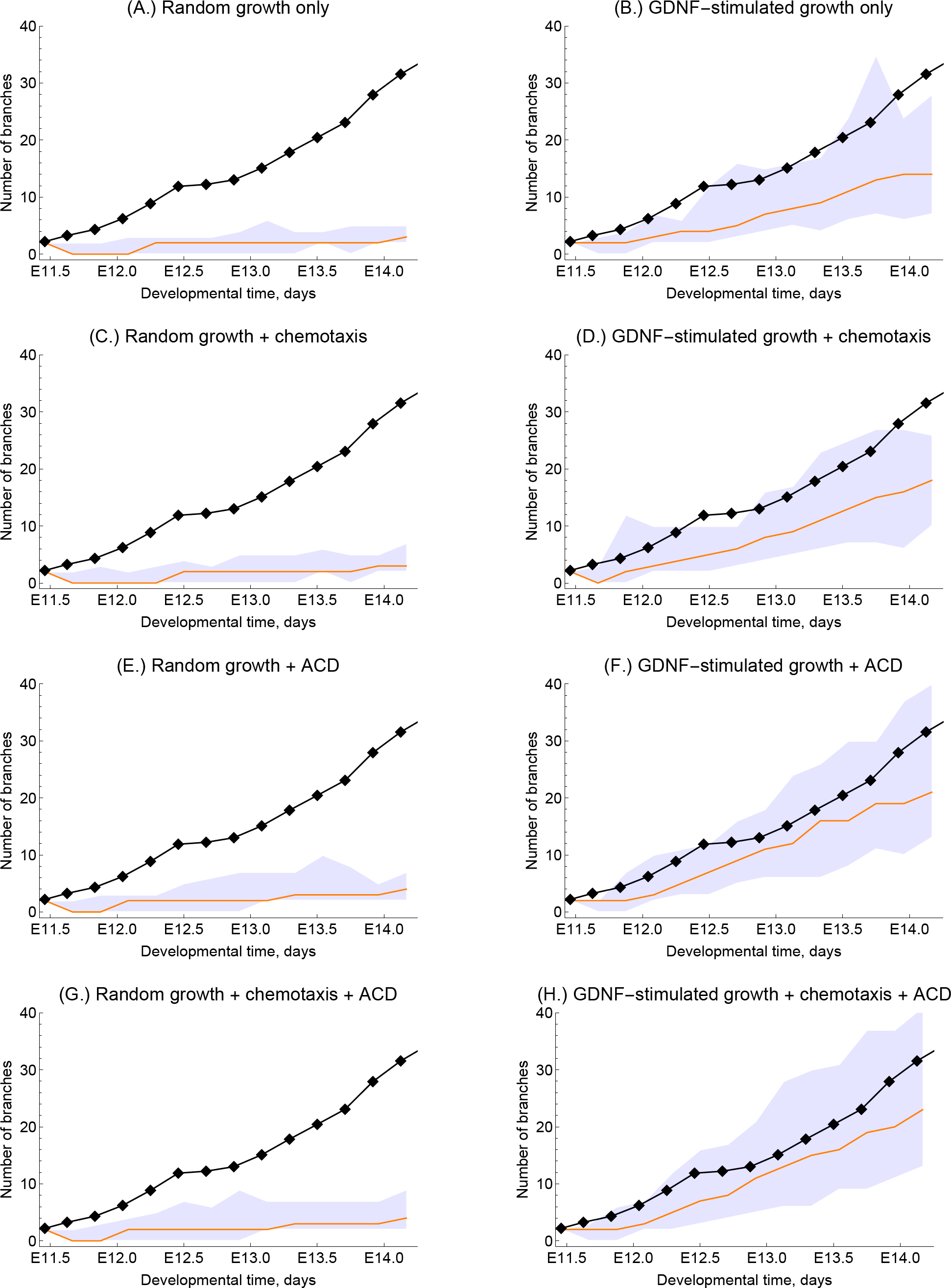
The effect of different GDNF signalling mechanisms on explant branching. (A.)-(H.) show simulation results with the indicated mechanisms implemented, and correspond to cell distributions shown in the panels of Fig. 5 with the same letters. In each panel the black line and points represent the evolution of branches from an explant experiment in Watanabe & Costantini (2004); the orange line represents the mean branching observed by model simulation (n= 200) and the shaded region indicates the 95% confidence interval. “ACD” indicates “anisotrophic cell division”. The parameter values used in each case are in Table S1.

**Figure 5.**
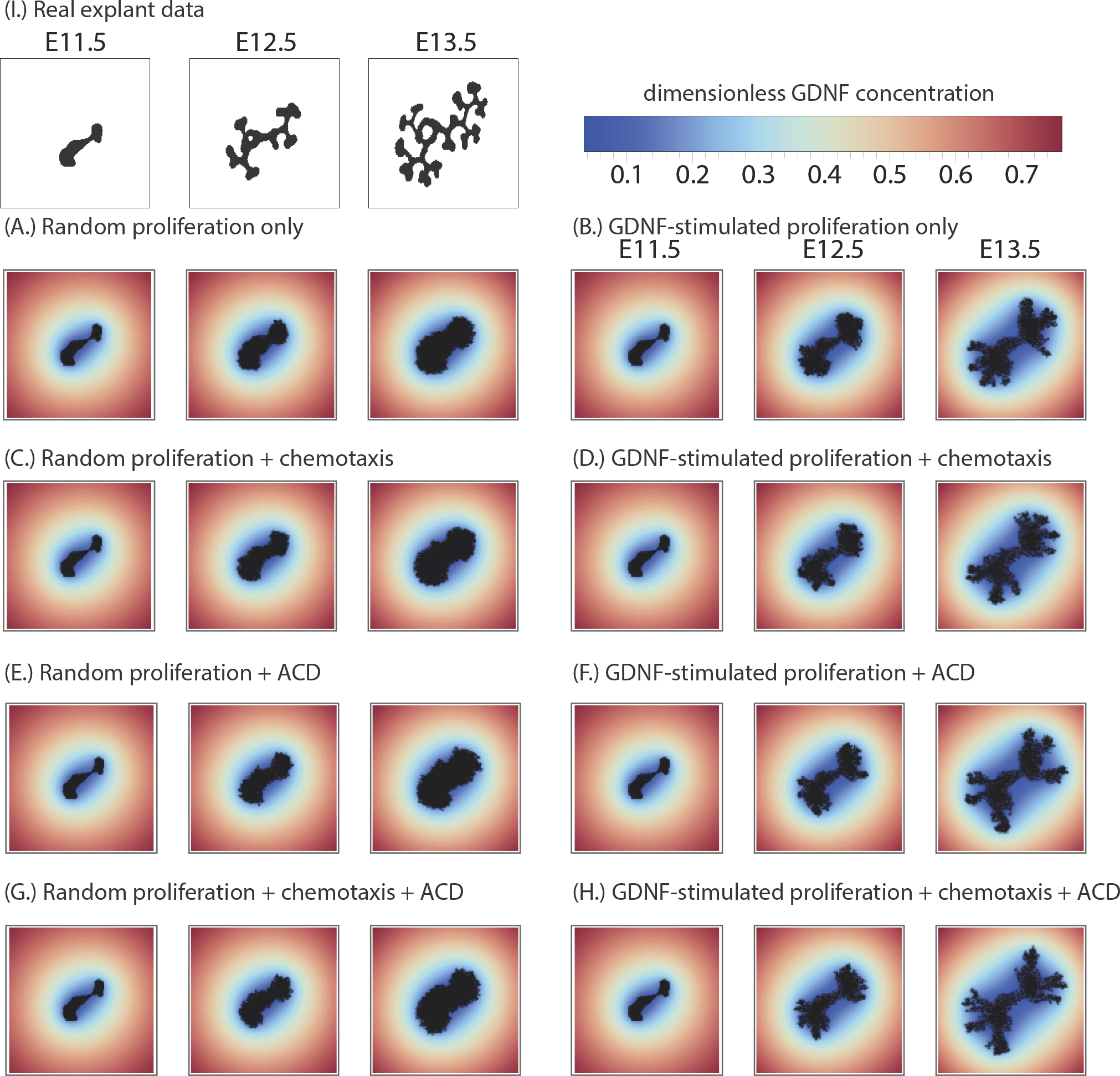
The effect of different GDNF signalling mechanisms on explant branching. (A.)-(H.) show simulation results with the indicated mechanisms implemented, and correspond to the panels in Fig. 4 with the same letters. The black mass is the epithelial cells, and the coloured shading shows the corresponding dimensionless distribution of GDNF. For comparison experimental from the first video in Watanabe & Costantini (2004) are presented in (I.). “ACD” indicates “anisotrophic cell division”. Note: the GDNF level at the edges is not *g_∞_* = 1 because the panels only show the central 140*×*140 grid points of the domain (itself of size 400*×*400). The parameter values used in each case are in Table S1.

**Figure 6.**
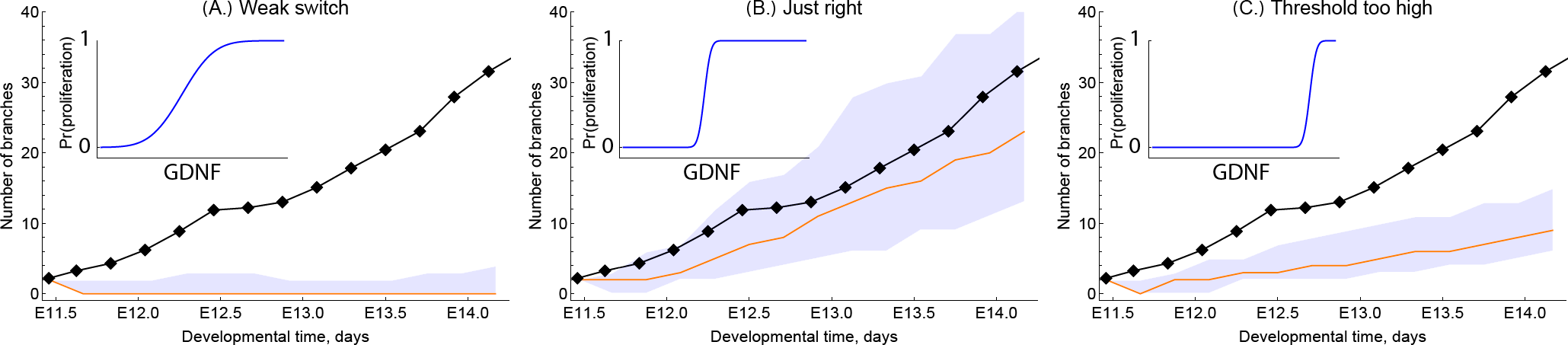
Series of simulation results showing how the shape of the GDNF-mediated proliferation switch influences the branching dynamics. For the results shown in each of the panels we ran simulations with *c*_1_ =− 25 so that in the absence of GDNF cells would not divide, and we vary *c*_2_ across each of the panels: for (A.) *c*_2_ = 400, (B.) *c*_2_ = 120 and (C.) *c*_2_ = 20. The inset panels show the location of the proliferation switch in GDNF-space (horizontal axis), against the probability of growth (vertical axis). In each panel the black line and points represent the evolution of branches from an explant experiment in Watanabe & Costantini (2004); the orange line represents the mean branching observed by model simulation (n= 200) and the shaded region indicates the 95% confidence interval. The parameter values used in each case are shown in Table S1.

**Table 1.** A summary of the effect of different GDNF signalling mechanisms on explant branching. Each row of the table corresponds to specific panels of Figs. 4 and 5, with corresponding parameter sets indicated in Table S1. In the middle three columns “-” indicates that the mechanism is inactive, “+” indicates that it is active. In the right-hand column “-” indicates no significant branching, “+” indicates modest branching and “++” indicates that branching that is consistent with experimental data.

| figure | GDNF-stimulated cell division | chemotaxis | anisotropic cell division (ACD) | branching |
| --- | --- | --- | --- | --- |
| 4,5A | - | - | - | - |
| 4,5B | + | - | - | + |
| 4,5C | - | + | - | - |
| 4,5D | + | + | - | + |
| 4,5E | - | - | + | - |
| 4,5F | + | - | + | ++ |
| 4,5G | - | + | + | - |
| 4,5H | + | + | + | ++ |

Model simulations reveal that when cell division is independent of GDNF, no branching occurs; the explant grows as an approximately circular mass (see Figs. 4, 5A). When only cell proliferation is regulated by GDNF, some branching occurs, although the number of branches is fewer than for explant growth (see Figs. 4, 5B). When chemotaxis and/or anisotropic cell division depend on GDNF although the rate of proliferation is independent of GDNF, the model does not exhibit branching (Figs. 4,5 C,E,G). Additionally the cumulative effect of GDNF-stimulated proliferation and chemotaxis on branching is no greater than proliferation stimulated by GDNF alone (Figs. 4, 5D). By contrast when proliferation and anisotropic cell division depend on GDNF, the number of branches observed along the branching trajectory increases such that simulation results are in good agreement with the experimentally observed branching patterns (Figs. 4, 5F). The best agreement with the experimental data is obtained when all three processes (i.e. proliferation, chemotaxis and anisotropic cell division) depend on GDNF levels (Figs. 4,5H). These results do not appear to depend strongly on the initial shape of the epithelium used to in model simulations. In particular simulations initialised using different experimental explant data yielded similar results (see Figs. S1, S2).

These simulation studies reveal several key messages. First, GDNF-stimulated proliferation can generate biologically realistic branching patterns. Second, what we term anisotropic cell division ameliorates the rate of branching when it is coupled to GDNF-stimulated growth. Finally, within the regions of parameter space studied, GDNF-controlled chemotaxis does not strongly affect the branching of explants. In order to determine whether the weak dependence on chemotaxis was due to an overly simplistic treatment of cell-cell interactions, we revised our CA model to ensure that epithelial cells remain attached to at least one neighbouring cell. However this change did not significantly alter the observed branching patterns (results not shown).

We also investigated how the shape of the GDNF-controlled switch that regulates cell proliferation affects branching (Fig. 6). When the slope of the switching function is gradual, the epithelium grows as a circular mass and no branching occurs (Fig. 6A). Similarly, when the threshold of the switch is high, the epithelial growth rate and branching rate are both too low (Fig. 6C). We therefore tune the threshold and slope of the GDNF-mediated switch (Fig. 6B) in order to maximise the amount of branching. Again we show in the Supplementary Materials how these results are robust to variation in the shape of the epithelial cell starting mass (see Figs. S3 and S4).

### 3.3 Branching is sensitive to the form of the GDNF proliferation switch

Having shown that our CA model can reproduce qualitative features associated with early kidney morphogenesis, we now study the dependence of branching characteristics on the parameters *c*_1_, *c*_2_, and *p_move_*; *c*_1_ and *c*_2_ jointly determine the sensitivity of cell proliferation to GDNF levels (see eqn. (6)), and *p_move_* is the probability of cell migration.

We performed 50,000 AABC simulations and accepted the top 5% (the 5% with overall smallest Euclidean distances from the summary statistics associated with the experimental data) to compose approximate posterior samples, as outlined in section 2.3. We use two summary statistics to compare simulations with data: the normalised area and the total number of branch points as calculated from medial axis skeletons of the epithelial cell mass.

In Fig. 7A we plot those trajectories that have been accepted as posterior samples. The marginal posterior distributions associated with each parameter are shown on the diagonal of Fig. 7B alongside the two-dimensional posterior joint density distributions for each pair of parameters. This reveals that the marginal posterior distribution for *c*_2_ deviates most from its prior, taking values between 40 and 280 with high probability; both *c*_1_ and *p_move_* deviate less from their prior, although *p_move_* shows deviation at high values where little or no growth occurs.

**Figure 7.**
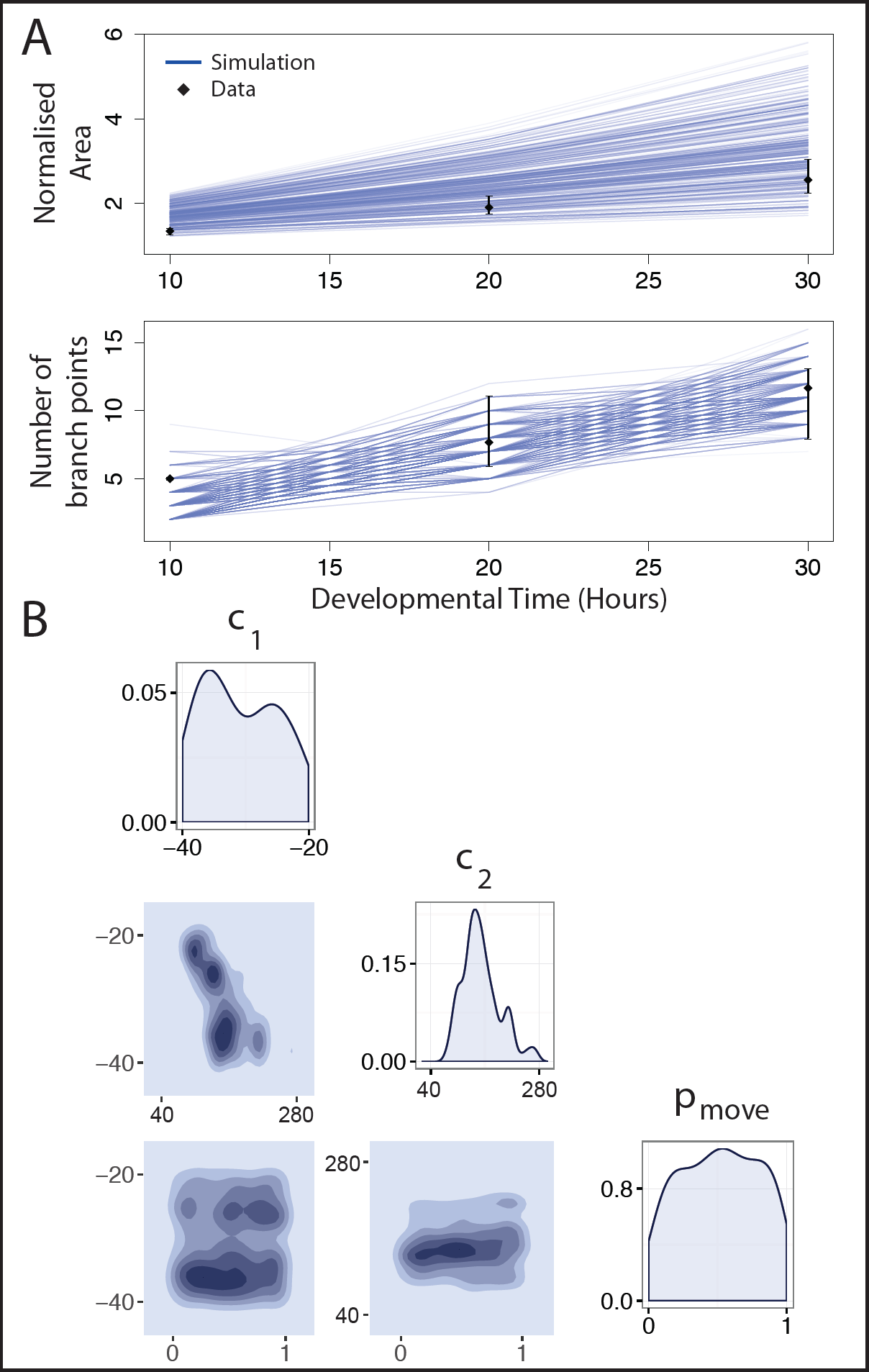
Approximate posteriors distributions for three model parameters as estimated by AABC. The parameters *c*_1_ and *c*_2_ affect the location and sensitivity of the GDNF-mediated growth switch, and *p_move_* determines the relative likelihood of cell movement rather than cell division (see “Materials and Methods”). (A) Accepted trajectories in the posterior simulated from the model and their comparison with the *ex vivo* data (black dots indicate means, and error bars show the range of the data). The experimental data are the means of summary statistics extracted from three explant experiment videos in Watanabe & Costantini (2004). (B) Posterior parameter distributions for single (on the diagonal) and joint pairs of parameters. See Table S1 for information about the parameter values used in the simulations.

The joint density plots in Fig. 7B reveal dependencies in parameter space. For the parameter pair (*c*_1_*, c*_2_), a negative correlation is observed; we note also that for low values of *c*_1_(i.e. *c*_1_ ∊ [−40 *, −*25]) *c*_2_ is tightly constrained. This dependence is expected since the two parameters jointly determine the sensitivity of the switch to GDNF. In particular, under a local non-dimensional concentration of GDNF of 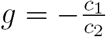 a cell with free neighbouring space divides with probability 0.5. This means that there are regions of (*c*_1_*, c*_2_) space (where *c*_1_ *∼ −kc*_2_) which result in practically identical GDNF switches. For the other two parameter pairs, the joint density plots highlight what has been termed “sloppiness” (Gutenkunst *et al.*, 2007): the explant branching phenotype is robust to changes in the values of *c*_1_ and *p_move_*. Parameter sloppiness can be symptomatic of several problems, including insufficient data, or a model that is overly complex for the available data. Additionally we cannot discount the possibility that our summary statistics are suboptimal, although we believe that the use of more detailed summary statistics may be excessive given the level of biological and mechanical realism present in our CA model.

## 4 Discussion

We have developed a new model to describe branching morphogenesis in the developing kidney. While there are many agent-based models of other branching systems (e.g. Fumoto *et al.* (2016); Iber & Menshykau (2013); Lindenmayer (1968); Merks & Koolwijk (2009); Schatten *et al.* (2007)) to our knowledge, this is the first agent-based model of kidney morphogenesis. A significant advantage of our approach is that unlike spatially-averaged compartment-based methods (e.g. Menshykau & Iber (2013); Zubkov *et al.* (2015)) it allows study of processes at the single-cell level. This spatial resolution may be particularly important in determining the thresholds in the concentration of growth factors at which branching occurs. The CA framework allows a cell-based description of tissue morphogenesis and facilitates the future addition of other biophysical mechanisms and the subcellular signalling pathways, as well as extension to include other cell types (e.g. mesenchymal cap cells, when considering kidney morphogenesis).

At present there is no consensus about whether diffusion-driven Turing patterns of GDNF coupled with GDNF-regulated proliferation mechanisms can explain branching, or whether other chemical and mechanical mechanisms are required. Our simulation results indicate that the GDNF-mediated proliferation may suffice to generate branching in this system. We find that to recapitulate the branching behaviour of the developing kidney, dynamic spatial patterning of epithelial cell proliferation had to be included in the model. This was the case across all the parameter sets that we considered, however there may be isolated regions in parameter space in which branching occurs via independent mechanisms not included in our model (e.g. due to mechanical cell-cell interactions). Simulation studies demonstrated that GDNF-stimulated tissue growth together with chemotaxis and anisotropic cell division provided the best fit to the explant branching data studied. In a recent experimental study, Riccio *et al.* (2016) studied the behaviour of tip cells across development, and concluded that GDNF signalling largely drives cell movement (rather than proliferation), and that this movement, in turn, drives branching. While GDNF-dependent chemotaxis may play a role later in kidney development (we focus here on E11.5 to E14), our results suggest that it may not contribute significantly during early branching of the epithelium, and supports a greater role for proliferation than than that identified by Riccio *et al.* (2016). It is, of course, possible that mesenchyme-derived factors other than GDNF provide this signal. Inference of the CA model on a three-dimensional subset of parameters revealed dependency between the parameters controlling the GDNF-stimulated cell proliferation switch (*c*_1_ and *c*_2_). Specifically this result indicated how cell proliferation should depend strongly on GDNF levels. However it remains to identify a biological mechanism that fulfils this criteria.

Whilst our model can generate explant patterns that mimic some aspects of the experimental data, we recognise that our simulation results often differ in the finer details. At present it is not clear whether these discrepancies are due to biological mechanisms that we have neglected, or due to the relatively simple nature of our model framework. For example these differences might be reduced if an off-lattice agent-based model, incorporating cell-cell contact forces, were used. We also note that the differences between our simulation results and the experimental data are highest for later developmental times, and we speculate that other mechanisms may be responsible for continued branching at these later time points. In particular our model does not explicitly include mesenchymal cells that produce growth factors which may influence branching at later developmental times. Another candidate mechanism is heterogeneity in RET expression amongst different epithelial cells (Shakya *et al.*, 2005). In particular, epithelial cells with differing levels of RET expression have been shown to compete for positions within branches (Riccio *et al.*, 2016), and contribute unequally to the tips of the developing kidney (Chi *et al.*, 2009). We also assumed that the time scale for GDNF diffusion is much slower than the time scale for its uptake/binding to RET receptors. However we cannot discount that future experiments may invalidate this assumption.

As important differences exist between the dynamics of branching as it occurs *ex vivo* and *in vivo* (Short *et al.*, 2014), caution should be exercised when extrapolating from the former to the latter. Here we focus on an *ex vivo* model, due to the availability of published image data. This system also lends itself more naturally to 2D modelling, which is less computationally expensive than 3D simulations. Scaling an ABM approach to 3D represents a significant computational hurdle, and it is possible that the different topology may require qualitatively different mechanisms for branching to occur. However our results are supported by the results of computational modelling by Clément & Mauroy (2014) who found that diffusion of growth factors was sufficient to generate realistic 3D branching patterns. Further our 2D model permits investigation of the dominant mechanisms in a simpler geometry, and presents an opportunity for a first model validation step before investigating 3D dynamics. As noted above our model omits certain details of the biology, for example, the (likely) presence of a growth factor-producing mesenchyme. This choice was dictated partly by the sparsity of the available experimental data. However since our simple model was able to reproduce key aspects of the explant branching we chose not to include further (uncertain) biological details.

The flexibility that CA models offer in their ability to describe spatiotemporal heterogeneities is not only an advantage but also a limitation, because the biological interpretation of each update rule is not always clear. Modelling cell proliferation and migration with biophysical and mechanical forces could improve the mechanistic understanding of the model, but comes at significant computational cost (see for example, Kim *et al.* (2007); Rejniak & Anderson (2011)).

To perform parameter estimation, we have implemented a version of AABC, an approximate Bayesian inference scheme that is ideally suited to models such as ours that are computationally expensive to simulate. With the ever-increasing resolution of spatiotemporal data and, concurrently, an increasing number of models developed to describe relevant biological phenomena, we propose that AABC may find useful applications to a range of problems in systems biology, outside the more typical population genetics applications for which it was developed (Beaumont *et al.*, 2002; Buzbas & Rosenberg, 2015; Csilléry *et al.*, 2010b).

Existing models of organ development have proposed alternative mechanisms for branching kidney organogenesis. In Zubkov *et al.* (2015), a spatially-averaged system of ODEs is proposed in which branching occurs at a specific cell ratio of epithelial (tip) and mesenchymal (cap) cells. In both Clément & Mauroy (2014) and Menshykau & Iber (2013), spatially-resolved models are developed and a growth-promoting ligand mechanism is proposed for branching; in Menshykau & Iber (2013), the authors show that this leads to a Turing-type mechanism through interaction of GDNF and the RET receptor. The model that we present is consistent with these results, but goes further by proposing cellular scale rules that, coupled to the influence of a ligand field, enable branching.

We recognise that other modelling frameworks have been used to simulate branching morphogenesis. These include phase field models (Hartmann & Miura, 2006; Ohta *et al.*, 1989) and Turing models (Kondo & Miura, 2010; Menshykau & Iber, 2013)), that yield similar conclusions to our CA approach. Even so, we believe that it is important to establish whether results are robust across different modelling approaches or specific to a particular modelling paradigm.

In conclusion we note that while branching morphogenesis is an old problem in mathematical biology (Murray *et al.*, 1983), many open questions remain to be addressed. Here, using an agent-based model that directly describes the cell–cell interactions that occur during organ development, we shed light on the processes involved in defining the structure of the kidney. In the future, we propose that more complex hybrid models that combine biophysical and experimentally-validated rules for the migration and proliferation of epithelial cells will lead to further advances in our understanding of kidney morphogenesis. Additionally we argue that future models of *in vivo* kidney development should include other cells types known to be involved in organogenesis, for example, mesenchymal cells.

## Acknowledgments

We thank Joe Pitt-Francis for useful advice on the image-processing of the explant data. We also thank the EPSRC for partial funding.

